# *Schistosoma Japonicum* infection in Treg-specific USP21 knock-out mice

**DOI:** 10.1101/2020.03.09.983502

**Authors:** Youxiang Zhang, Guina Xu, Baoxin Zhang, Shan Zhang, Yangyang Li, Qing Huang, Simin Chen, Fansheng Zeng, Bin Li, Zhiqiang Qin, Zuping Zhang

## Abstract

USP21, an E3 de-ubiquitin enzyme playing vital roles in physiological activities, is important for Treg cells to maintain immune homeostasis and control immune tolerance. To understand its diverse functions and potential mechanism is essential for disease development. We, using the USP21 gene-conditional knock-out mice model of *Schistosoma Japonicum* infection, found more cercariae developed into adults, and more eggs deposited in the liver in KO mice. However, immunohistochemistry showed the degree and the area of egg granuloma and liver fibrosis were both reduced. This suggested knock-out USP21 did affect the immunoregulation between schistosomes and the host. In KO mice the content of IFN-gamma and IL-4, and the expression of anti-SEA IgG and anti-SWAP IgG both increased in the liver, spleen and blood by flow cytometry, while the content of IL-10, lL-17A, IL-23, IL-9 and the expression of USP21 and anti-SEA IgM decreased. This indicated USP21-knockout-Tregs promoted both Th1-type and Th2-type immunity and inhibited other immunities during schistosomes infection, which disordered the host immunity. This study revealed the immunomodulatory of USP21 and preliminarily suggests it might be essential to regulate the complex immune network between the host and schistosomes. USP21 provides a new possible target for schistosomiasis treatment in the future.

**Author summary:** Schistosomiasis is a common neglected tropical disease that affects more than 230 million people worldwide. Therefore, the study on the mechanism of immune interaction between schistosomas and the host is not only helpful for the understanding of immune homeostasis, but also helpful for the further development of the treatment of schistosomiasis. Ubiquitin Specific Protease 21(USP21) has been shown to be involved in the regulation of many biological processes, such as maintaining immune homeostasis and regulating cell growth. Here, we observed that the specific deletion of USP21 led to the decrease of mice’s ability to resist schistosomes infection and promoted the survival of schistosomes. It was also proved that unstable regulatory T cells would produce polarization phenomenon and promote differentiation to helper T cells, which would lead to disorder of immune response in mice. However, this process reduced the serious immune pathological damage caused by egg granuloma. Our findings reveal that USP21 may be an important molecule regulating immune interaction between *Schistosoma japonicum* and the host.

## Introduction

Schistosomiasis is an acute and chronic parasitic disease caused by blood flukes (trematode worms) of the genus *Schistosoma*. According to the WHO Report 2017, Schistosomiasis transmission was reported from 78 countries, and at least 220.8 million people required preventive treatment in 2017. The estimates of death due to Schistosomiasis varies between 24 072 and 200 000 globally per year. In 2000, WHO estimated the annual death at 200 000 globally[1]. People become infected when larval forms of the parasite–released by freshwater snails– penetrate the skin during contact with infested water. In the body, the larvae develop into adult schistosomes. Adult worms live in the blood vessels where the females release eggs. Some of the eggs are passed out of the body in the feces or urine to continue the parasite’s lifecycle. Others become trapped in body tissues, causing immune reactions and progressive damage to organs[2]

The immune systems of infected hosts have several life cycle stages of the parasite that it must confront: penetrating cercariae, migrating schistosomula, adult worms and the eggs produced by adult worm pairs. The developing stages of the parasite can express hundreds of antigenic moieties, many of which stimulate strong humoral and cellular immune responses[3, 4]. Some of these responses continue to increase during chronic infection, and others are strongly down-regulated [5-7]. Observations in murine experimental infection models have shown the mechanisms governing the development and regulation of the pathogenic immune response in Schistosomiasis [8-11]. The mechanisms of immunomodulation include IL-10, T regulatory cells, B cells, antibodies, and T cell anergy[12-15]. Besides, several studies have suggested that Th17 cells are involved in the pathogenesis of both *S. mansoni* and *S. japonica* [16-18].

Treg cells (Tregs) have the immunosuppressive capacity, essential for maintaining immune homeostasis and controlling immune tolerance. It has been shown that inducing of Treg cells can inhibit the development of granuloma pathology [19-24]. FOXP3 is a crucial factor in the phenotype of Treg cells to obtain immunosuppressive activity. Unstable FOXP3 will keep the Treg cell phenotype stable, may bias different T helper cell-like phenotypes in different inflammatory conditions[20, 21, 25, 26]. Further, the conditional knock-out mouse model found that USP21 stabilizes the expression of FOXP3 protein by deubiquitination in regulatory T cells, thereby regulating the function of Treg cells. A Th1-like sputum cell response, as expressed in cells of the USP21-deficient Treg, preferentially becomes a Th2 cell-like phenotype in a host with a severe Th2-type disorder, whereas, in arthritic conditions, regulatory T cells may lose FOXP3 expression and transform into Th17 cell-like cells [23, 27-30]. More importantly, the three (FOXP3 and GATA3, USP21) are closely related to regulatory T cells. In regulatory T cells, FOXP3 binds to the promoter region of USP21 and activates its transcription, while USP21 interacts with GATA3 and deubiquitinates it, inhibiting its degradation by the proteasome to maintain its protein stability.

Moreover, GATA3 can regulate the function of T cells in the inflammatory response by stabilizing the expression of FOXP3. Thus, in the regulatory T cells, the above three form a positive feedback pathway. Further, conditional knock-out mouse models found that USP21 regulated the expression of FOXP3 protein by deubiquitination in regulatory T cells, thereby regulating the function of Treg cells [26, 31]. In the antiviral reaction, USP21 can bind to and deubiquitinate RIG-I in the cytoplasm to play an immunomodulatory role, or hydrolyze the K27/63-linked polyubiquitin chain on STING to negatively regulate DNA virus-induced type I interferon[32, 33].

Although regulatory T cells may play an essential role in the immune regulation of schistosomes infection, the molecular mechanisms are not yet precisely defined. Therefore, this study aims to explore the mechanism of action of USP21 in the immune regulation of schistosomes infection by using the USP21 gene knock-out mice model. It will describe the conditional knock-out of USP21 mice infected with related immune molecules in *Schistosoma japonicum* and further study the molecular mechanism of USP21’s immune response to *Schistosoma japonicum.*

## Results

### Depletion of USP21 in Treg cells weakens resistance to *Schistosoma japonicum* in infected mice

In order to observe the difference between the KO mice and the WT mice, both were infected with *Schistosoma japonicum*, and the adults were recovered (according to the steps shown above), the number of females and male were combined, and the total adults were counted. We weighed the liver and spleen of the mice and observed the changes in their color, shape and texture. (Fig 1B and Fig 1D). The number of combined adults and total adults recovered from KO mice was significantly more than that from WT mice (Fig 1A). However, no effect on the development of cercariae into male and female adults was found.

**FIGURE 1.**
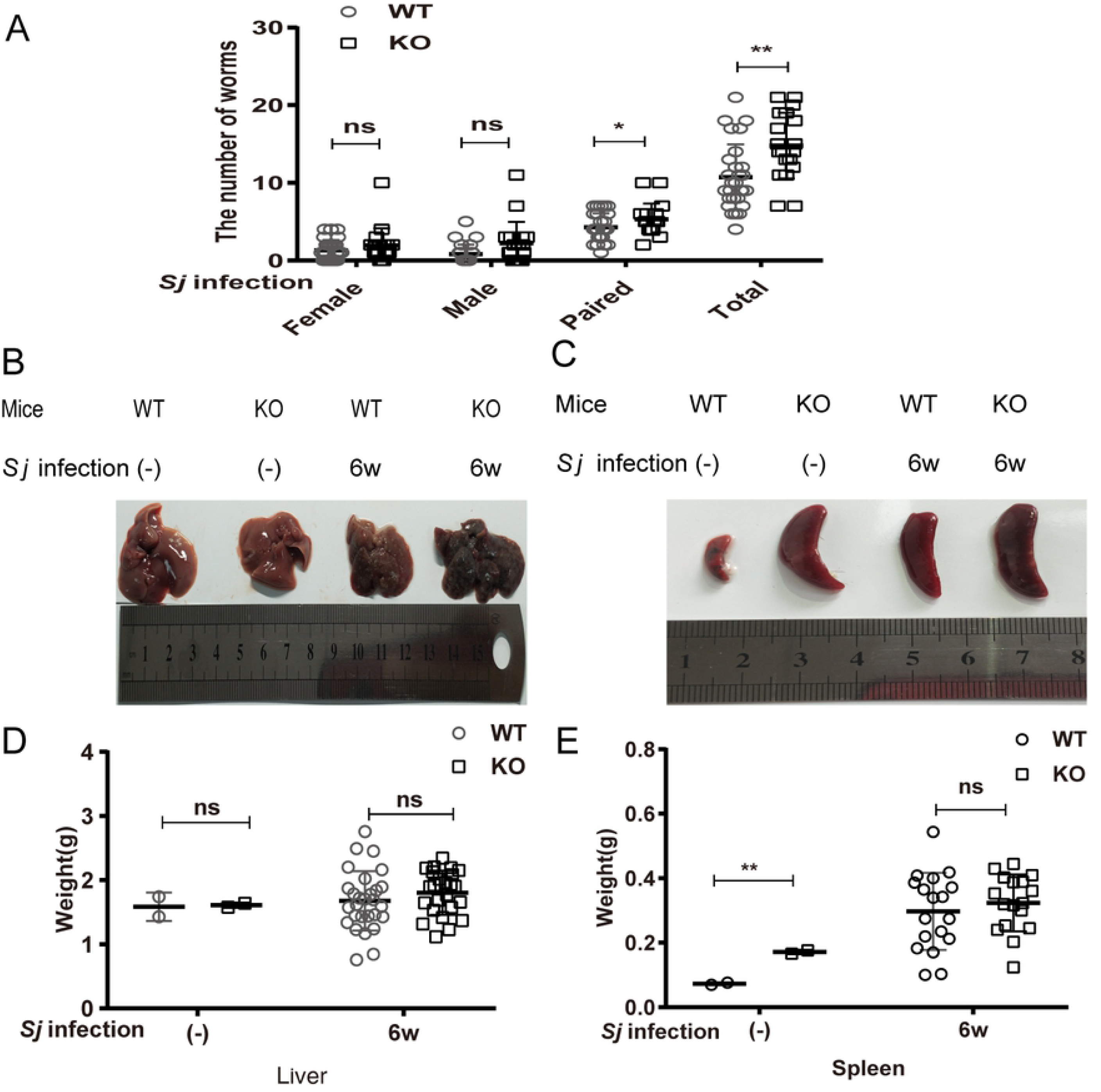
Depletion of USP21 in Treg cells weakens resistance to *Schistosoma japonicum* in infected mice. (**A-E**) FOXP3^Cre^(WT) and USP21^fl/fl^FOXP3^Cre^(KO) mice(n=26 ±2/group) were infected with 25±2 cercariaes through the abdomen, and were sacrificed at 6-7 weeks after infection. Liver and spleen tissues were collected and the adults were recovered. (**A**) Comparision of the number of females, males, hugs and adults recovered from WT and KO infected INF groups (n=mean ±SD), ^*^*p*=0.01in the total number of adults,and ^**^*p*=0.036 in total number of hugs.(**B**) Representative images of the liver of WT and KO, including NC and INF group. (**C**) Liver weight of WT and KO including NC and INF group(weight=mean ±SD), *p*>0.05 was considered no statistically significant between the normal control group and the infected group.. (**D**) Representative images of the spleen of WT and KO, including NC and INF group. (**E**) Spleen weight of WT and KO including NC and INF group,(weight=mean ±SD), ^**^*p*=0.007 between the normal control group, p>0.05 represented no statistical significance in the infected group.

### Effect of USP21 on *Schistosoma Japonicum* eggs in infectious KO mice

A part of the liver was taken for HE staining from the mice 42 days after infection in order to observe the pathological changes of egg granuloma. We analyzed the granuloma area and egg number in KO mice and WT mice using appropriate software. The results showed that the inflammatory cells of the egg granuloma of the KO mice were less than that of WT mice, and the color of HE staining was lighter in KO mice (Fig 2B). The area of liver egg granuloma in KO mice was significantly smaller than that in WT mice (Fig 2C), but the number of eggs in KO mice was significantly higher than that in WT mice (Fig 2A).

**FIGURE 2.**
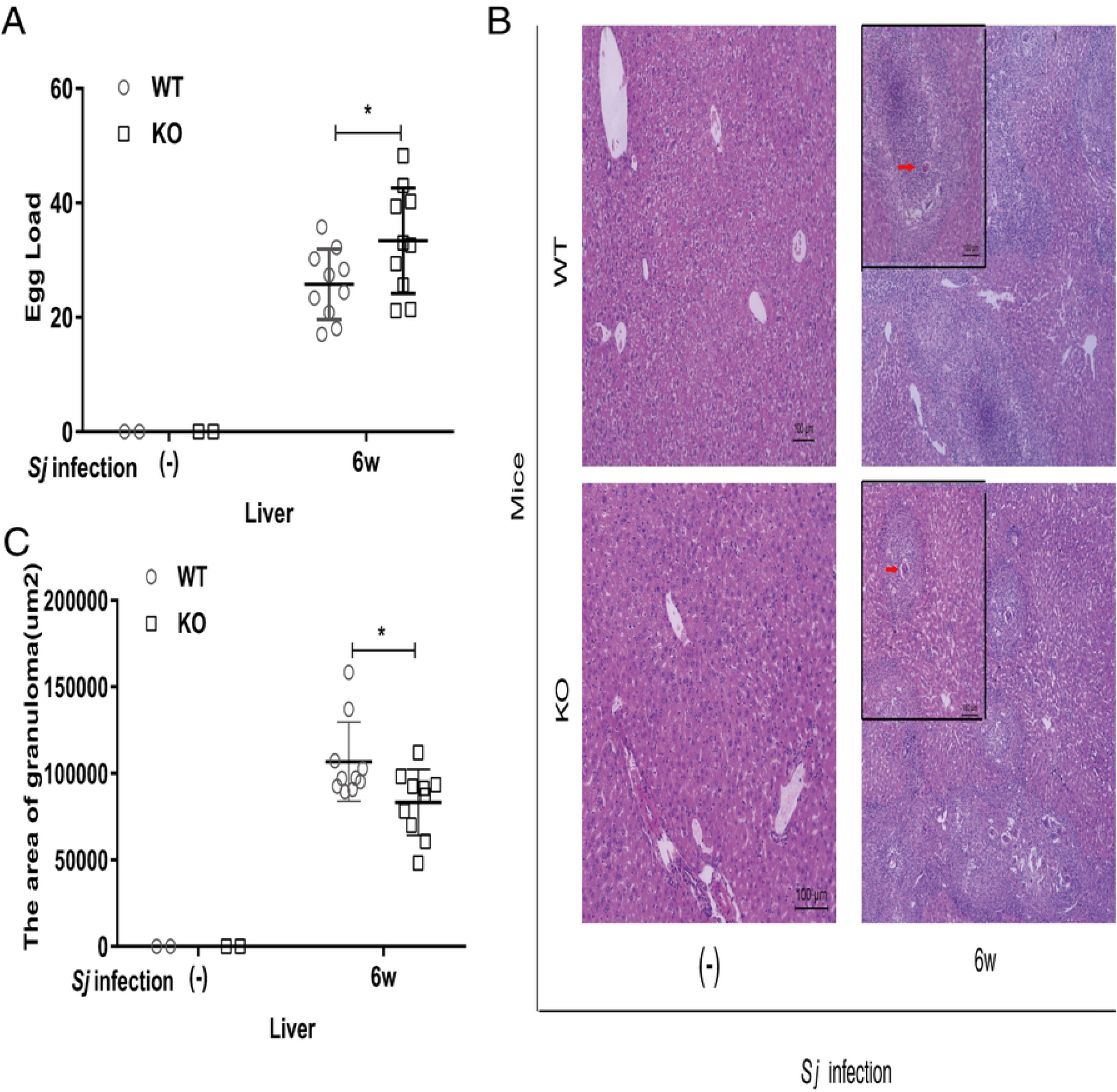
Effect of USP21 on *Schistosoma Japonicum* eggs in infectious KO mice. (**A-C**) The liver was taken for HE staining and the number of eggs was counted under the microscope on the 42nd day of the schistosomiasis infected mouse(n=10/group). (**A**) The statistical graph of liver eggs in `WT-INF` group and `KO-INF` group (the data expressed as mean ±SD, ^*^*p*=0.043). (**B**) Representative images of liver H&E staining in the WT-INF and KO-INF groups (original magnification: ×100, insertion: ×40).The arrows indicated the granuloma. (**C**) Comparison of liver egg granuloma area between WT-INF and KO-INF group. The data was expressed as mean ±SD,^*^*p*=0.022.

### Changes of liver Fibrosis in USP21^fl/fl^FOXP3^cre^ mice infected with *Schistosoma japonicum*

The proliferation of collagen fibers was observed by Masson trichromatic staining, which reflected the severity of hepatic fibrosis (Fig 3A). The mRNA expression of α-SMA, collagen-I and collagen-III was detected by RT-PCR and the protein expression of them were analyzed by Western blot. The area of collagen fiber in KO mice was smaller, and the color was lighter than that in WT mice, which was statistically significant (Fig 3B). The content of the α-SMA, collagen-I and collagen-III in the KO mice was significantly less than that of the WT mice at mRNA and protein levels (Fig 3C and Fig 3D).

**FIGURE 3.**
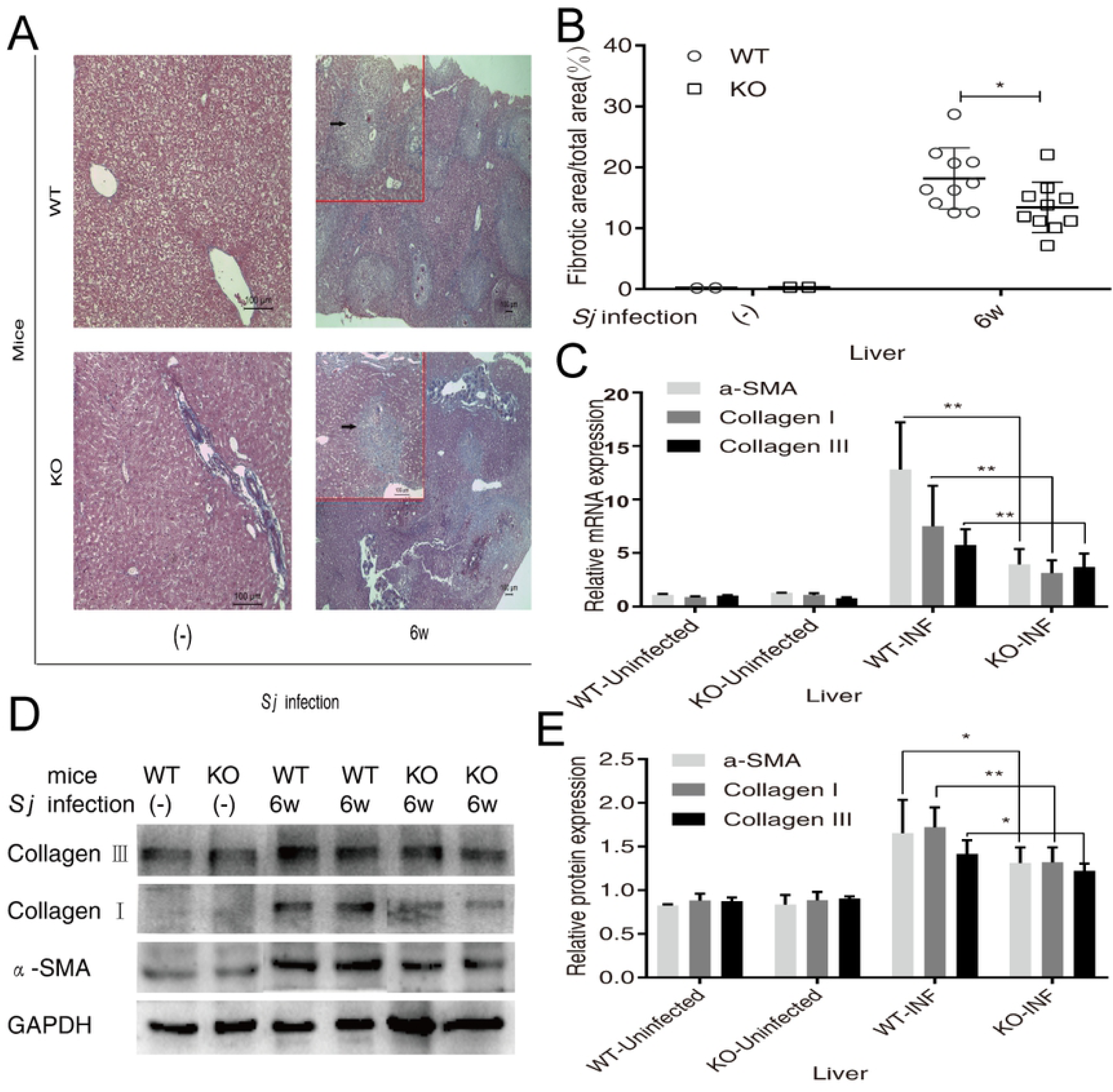
Changes of liver Fibrosis in USP21^fl/fl^FOXP3^cre^ mice infected with *Schistosoma japonicum*. (**A-E**) The liver was taken and masson stained on the 42nd day of the schistosoma infected mouse(n=10/goup).(**A**)Representative images of collagen deposition in WT-INF and KO-INF mice (original magnification: ×100) were obtained. The blue areas were collagen granules.(**B**) Comparison of the percentage of hepatic fibrosis area between WT-INF and KO-INF group. The data was expressed as mean ±SD,^*^*p*=0.039.(**C**)The mRNA level comparison of α-SMA, collagen-I and collagen-III `in` WT and KO including NC and INF group, the data representation for mean±SD. The mRNA level comparison of α-SMA, collagen-I and collagen-III between infectious groups, the data expresed as mean±SD. In infection group`α` -SMA:**p<0.001; collagen-I :^**^*p*=0.008, collagen-III :^**^*p*=0.006. (**D**) Representative images of SMA, collagen-I and collagen-III protein expression by western blot in WT and KO INF including NC and INF group(**E**) Comparison of protein expression between WT and KO including NC and INF group, the data expressed as mean ±SD. In infection group`α` -SMA : ^*^*p*=0.034, collagen-I:^**^*p*=0.006, collagen - III:^*^*p*=0.03.

### Changes of spleen immune in USP21^fl/fl^FOXP3^cre^ mice infected with *Schistosoma Japonicum*

To understand the difference of spleen immune cells between KO mice and WT mice after infection with *Schistosoma japonicum*, the spleen cells of the two groups, including normal control group, were collected, isolated and then cultured in vitro to detect and analyze the number of Treg cells and the percentage of CD4^+^CD25^+^FOXP3^high^ cells in CD4^+^ cells. The RT-PCR detected the changes of T cell types in the detection of relative mRNA expression of IL-10, IL-17, FOXP3, USP21, and by Multiplex Fluorescent Microsphere Immunoassay in the detection of the content of IFN-gamma, IL-4, IL-10, IL-17A, IL-23, IL-9 in splenocytes. mRNA levels of IL-10, IL-17, FOXP3 and USP21 in KO mice were less than that of WT mice (Fig 4C). The Flow Cytometry results were shown in Fig 4A and Fig 4B, and Splenic Cell Culture Cytokine results were showed as Fig 4D.

**FIGURE 4.**
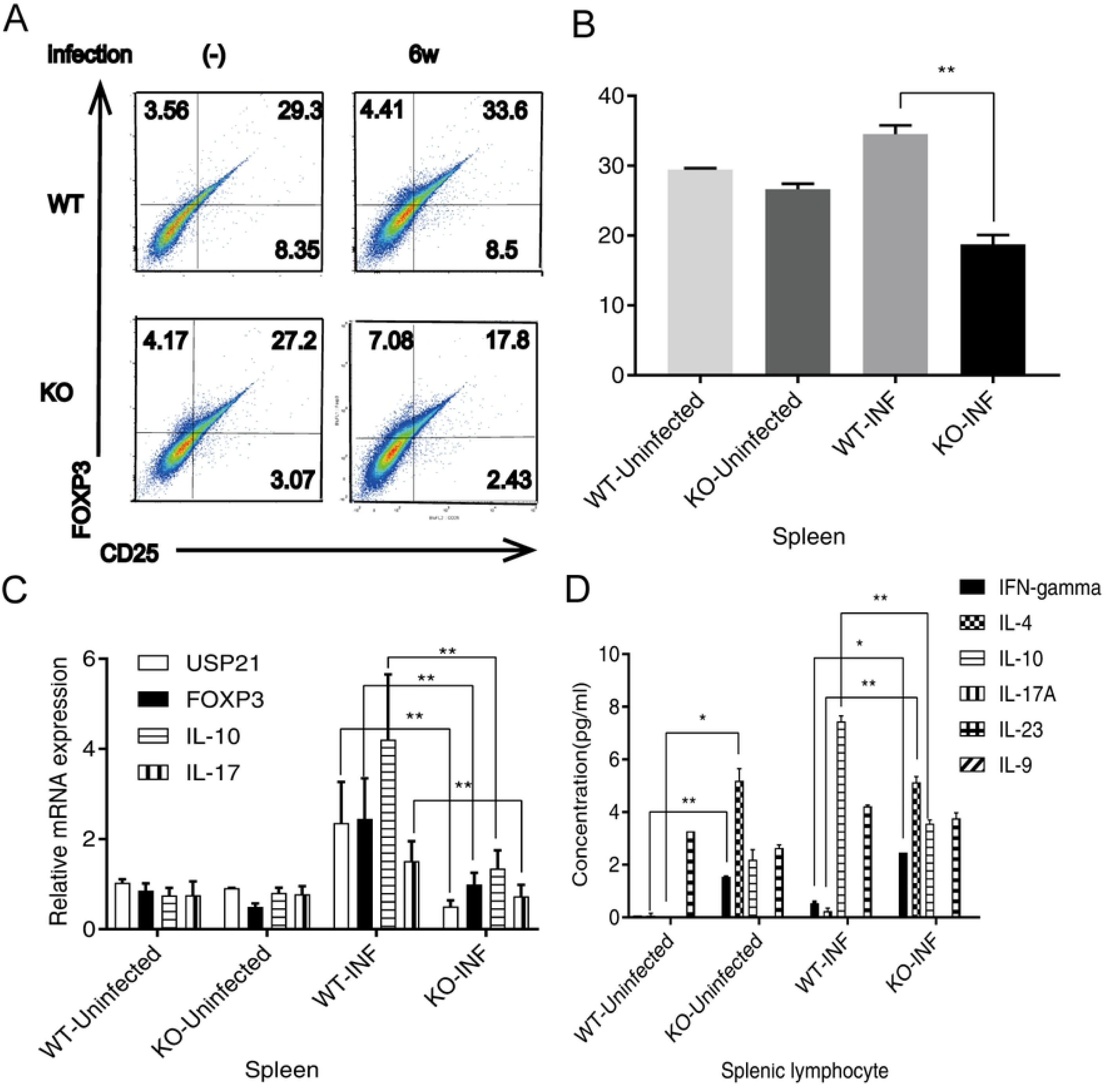
Changes of spleen immune in USP21^fl/fl^FOXP3^cre^ mice infected with *Schistosoma Japonicum*. (**A-D**)Spleen was collected and splenic cells were cultured in vitro on the 42^nd^ day in schistosomiasis infected mice(n=6/group).(**A**) CD4^+^CD25^+^FOXP3^high^ cells were determined by flow cytometry in WT and KO mice infected with schistosoma japonicum after 42 days, including NC and INF group,. (**B**) The comparison of the portion of CD4^+^CD25^+^FOXP3^high^ cells in CD4^+^T cells in INF of WT and KO, including NC and INF group. The data was expressed as mean ±SD, infection groups:^**^*p*=0.007.(**C**)The mRNA levels of FOXP3, IL-10, IL-17 and USP21 in WT and KO including NC and INF group, were compared, and the data was expressed as mean ±SD. Between infection group FOXP3 ^**^*p*=0.001,IL-10 ^**^*p*<0.001,IL-17 ^**^*p*<0.001, USP21 ^**^*p*<0.001.(**D**)The contents of IFN-gamma, IL-4, IL-10, IL-17A, IL-23 and IL-9 in cultured spleen cells of WT and KO including NC and INF group were compared, and the data was expressed as mean ±SD. In NC group IFN-gamma ^**^*p*=0.002, IL-4 ^*^*p*=0.002; between infection groups IFN-gamma ^*^*p*=0.013, IL-4 ^**^*p*=0.005,IL-10^**^*p*=0.004.

### Liver immunity in USP21^fl/fl^FOXP3^cre^ mice infected with *Schistosoma Japonicum*

In order to study whether the immune status of the liver was consistent with the condition of the spleen, the study detected the mRNA expression of IL-10, IL-17 and FOXP3 in the liver. We found that the mRNA expression of IL-10, IL-17 and FOXP3 in KO mice was significantly lower than that in WT mice (Fig 5D). The relatively large difference noted might be related to the small proportion of T cells in the liver. The mRNA and protein expression levels (Fig 5A) of USP21 in the liver were also lower in KO mice than that in WT mice (Fig 5B and Fig 5C).

**FIGURE 5.**
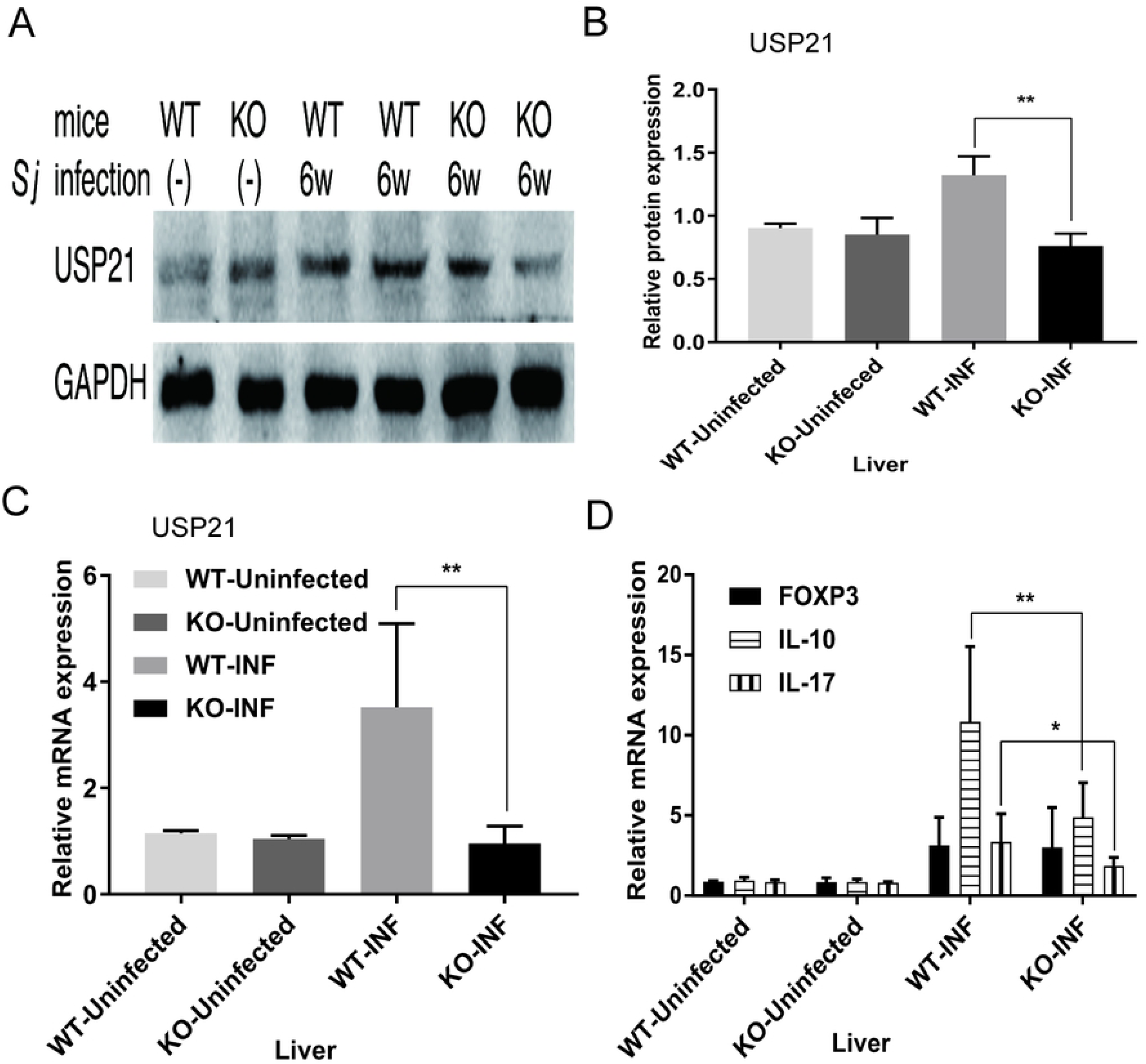
Liver immunity in USP21^fl/fl^FOXP3^cre^ mice infected with *Schistosoma Japonicum*. (**A**) Representative maps of USP21 protein expression were determined by Western blot in WT and KO including NC and INF group(n=6/group).(**B**)Protein expression of WT and KO, including NC and INF group(n=6/group), was compared. The data was expressed as mean ±SD, and the comparison between the infection groups was^**^*p*<0.001.(**C**)Comparison of USP21 mRNA levels between WT and KO, including NC and INF group(n=6/group), was presented as mean ±SD, ^**^*p*=0.001 between the infection groups.(**D**)The mRNA levels of FOXP3,IL-10, and IL-17 in WT and KO, including NC group and INF group(n=6/group), were compared, and the data was expressed as mean ±SD, between the infection groups IL-10 ^**^*p*=0.003, IL-17 ^*^*p*=0.026

### Specific antibody response in USP21^fl/fl^FOXP3^Cre^ mice infected with *Schistosoma japonicum*

To better understand the specific antibody response of USP21 ^fl/fl^FOXP3^Cre^ mice to *Schistosoma japonicum*, Serum samples from different stages were collected to measure the content of anti-SEA and anti-SWAP IgG/IgM antibodies from KO and WT mice in different infectious stages. The results were shown as follows: anti-SEA IgG (Fig 6A), anti-SEA IgM (Fig 6B), anti-SWAP IgG (Fig 6C), and anti-SWAP IgM (Fig 6D).

**FIGURE 6.**
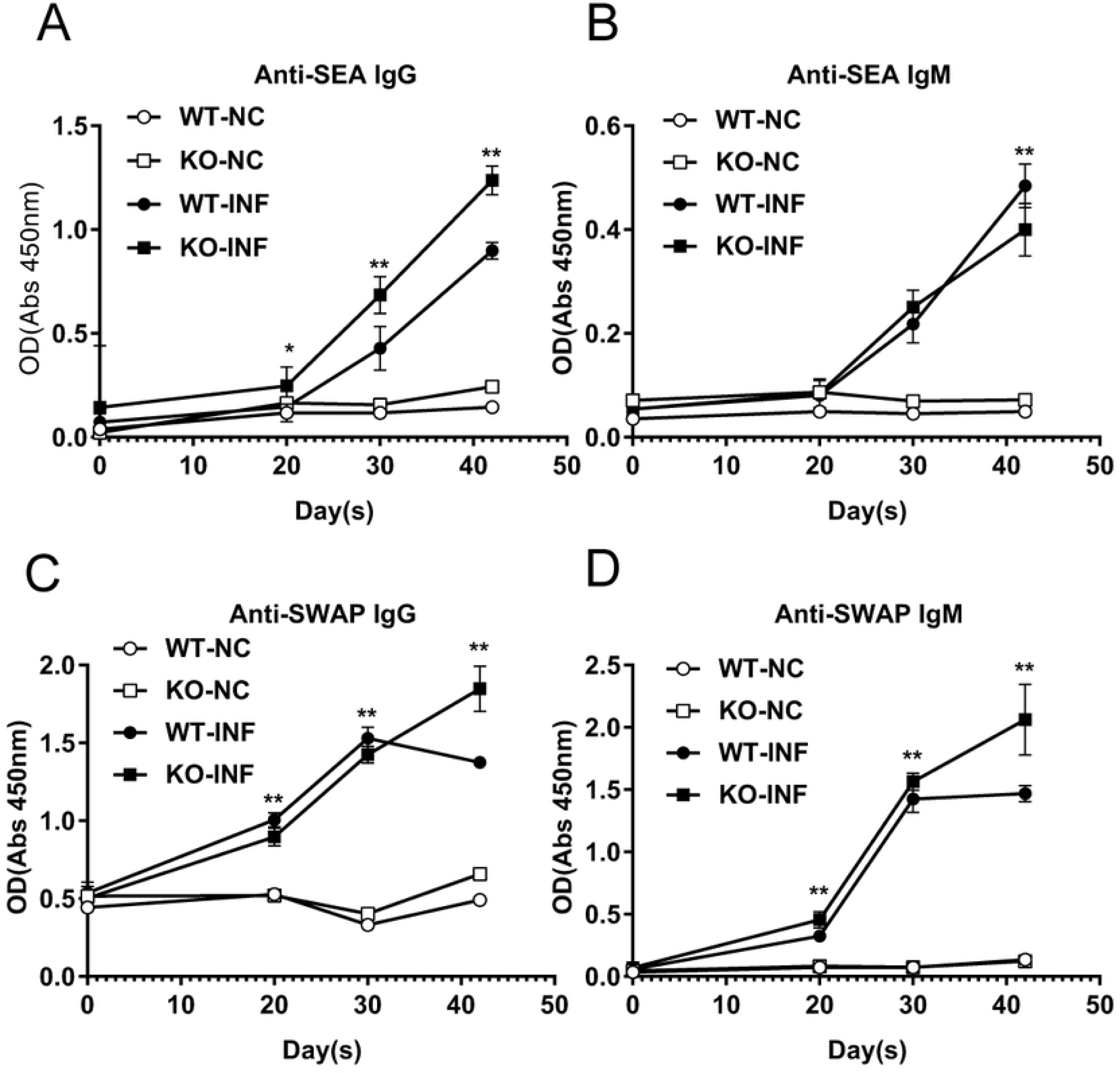
Specific antibody response in USP21^fl/fl^FOXP3^Cre^ mice infected with *Schistosoma japonicum*. (**A-D**)The serum from WT and KO including NC and INF group(n=26 ±2/group) at different stages (uninfected, day 20, day 30, day 42). (**A**) The changes of anti-SEA IgG content were compared, and the data was expressed as mean ±SD. Between infection group^*^*p*=0.01 on day 20, ^**^*p*<0.001 on day 30, and^**^*p*<0.001 on day 42.(**B**) Comparison of anti-SEA IgM content was made, the data was expressed as mean ±SD, and ^**^*p*=0.002 on day 42 between the infection groups.(**C**)Comparison of anti-SWAP IgG contents was taken to be expressed as mean ±SD, between infection groups^**^*p*<0.001 on day 20, ^**^*p*=0.003 on day 30, and ^**^*p*<0.001 on day 42.(**D**)The content of anti-SWAP IgM was compared, and the data was expressed as mean ±SD. Between infection groups ^**^*p*<0.001 on day 20, ^**^*p*=0.004 on day 30,^**^*p*<0.001 on day 42.

### Serum cytokines in USP21^fl/fl^FOXP3^Cre^ mice infected with *Schistosoma japonicum*

To intensively study on how Treg cells lacking USP21 affect mice resistance to *Schistosoma japonicum*, we tested the content of IFN-gamma, IL-4, IL-10, IL-17A, IL-23 and IL-9 in serum samples and PBMC from different infectious stages mice, including KO and WT mice. The results were shown as follows: comparison of the changes in the trend of the two groups of cytokines, including IFN-gamma, IL-4, IL-10, IL-17A, IL-23 and IL-9(Fig 7B), and comparison of cytokines in cell culture in vitro (Fig 7A).

**FIGURE 7.**
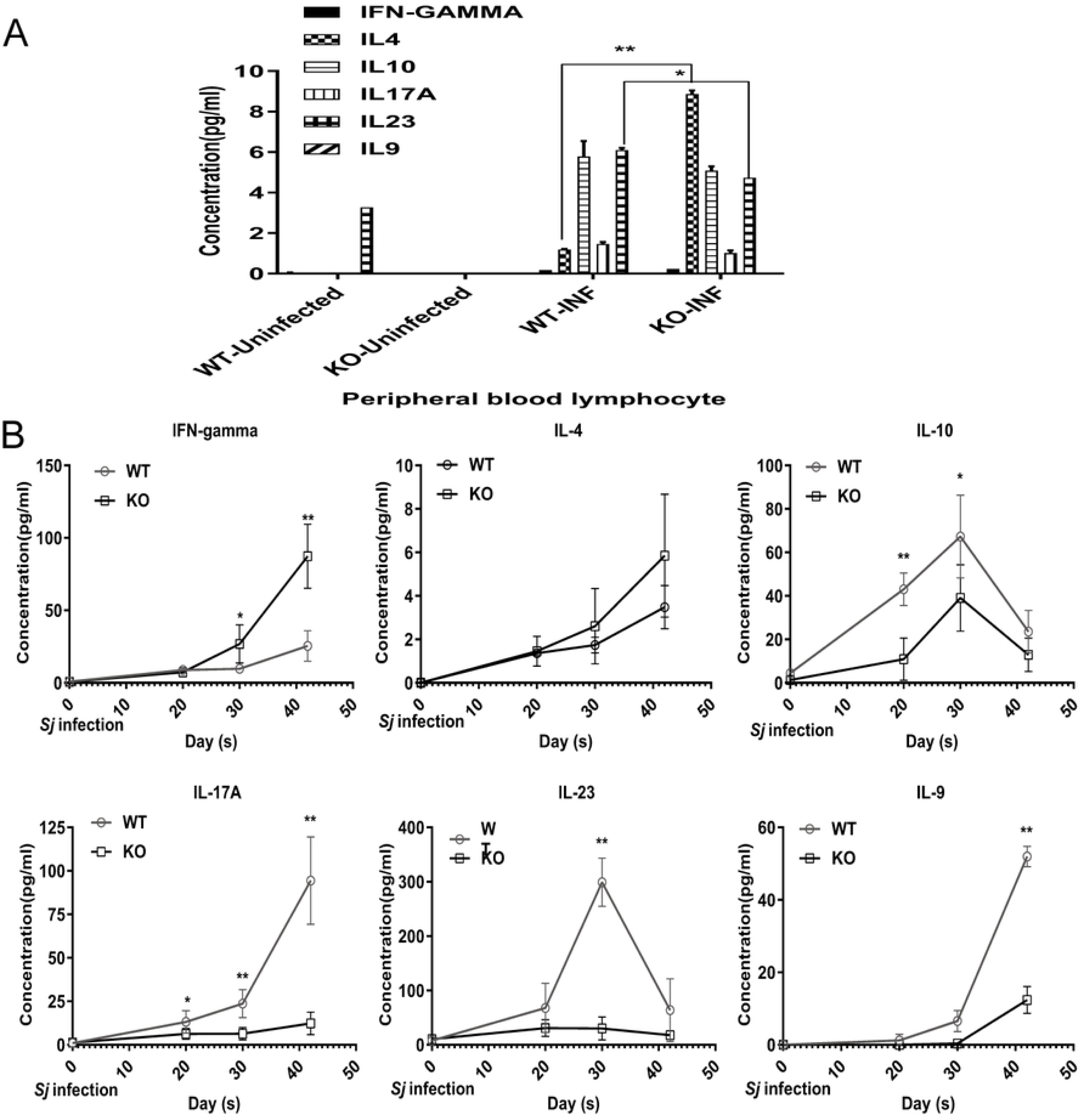
Serum cytokines in USP21^fl/fl^FOXP3^Cre^ mice infected with *Schistosoma japonicum*. (**A**)The contents of IFN-gamma, IL-4, IL-10, IL-17A, IL-23 and IL-9 in isolated peripheral blood lymphocyte cultures on day 42 of WT and KO, including NC and INF group(n=10/group), were compared, and the data was expressed as mean ±SD, IL-4 ^**^*p*=0.008,IL-23 ^*^*p*=0.035between the infection groups.(**B**)Contents of serum IFN-gamma, IL-4, IL-10,IL-17A, IL-23 and IL-9 was compared in INF of WT and KO infection groups(n=26±2/group)at different periods (uninfected, 20th day of infection, 30th day of infection and 42nd day of infection), and the data was expressed as mean ±SD. IFN-γ ^*^*p*=0.024 on day 20 and ^**^*p*<0.001 on day 42; IL-10 ^**^*p*<0.001 on day 20 and ^*^*p*=0.017 on day 30; IL-17A ^*^*p*=0.026 on day 20, ^*^*p*<0.001 on day 30 and ^**^*p*<0.001 on day 42;IL-23 ^**^*p*<0.001 on day 30 ; IL-9 ^**^*p*<0.001 on day 42.

## Discussion

USP21 is a member of the ubiquitin-specific proteolytic enzyme (USP) family and plays different biological functions. For example, USP21 regulated the gene expression of hepatocytes during liver regeneration by catalyzing the hydrolysis of ubH2A[34]. In inflammation, USP21 negatively regulated RIPK1 to inhibit the activity in the downstream of TNF-αR1[31], and USP21-mediated deubiquitination of IL-33 promoted the transcription of NF-κB p65[35]. Also, USP21 could bind to and deubiquitinate RIG-I in the cytoplasm to play an immunomodulatory role in antiviral reactions[32]. The latest studies have also shown that USP21 could regulate the expression of the cell cycle, proliferation, and craniofacial development related factors by deubiquitinating FOXM1 and Goosecoid (GSC)[36, 37]. We have previously used USP21 knockout mice model to explore their biological role in the growth and development of mice, and the differentiation and growth of lymphocytes and hematopoietic stem cells[38]. It is known that USP21 knock-out elderly mice showed spontaneous T cell activation and splenomegaly, and that the autoimmune system disorder was caused by Th1-like Tregs induced by the unstable expression of FOXP3 in USP21 deficient Treg mice model [26].

All stages of the schistosome life cycle inside the mammalian host elicit immune responses, but only the eggs are the target of a vigorous granulomatous inflammatory reaction. The immunopathological damage caused by *Schistosoma japonicum* is the result of promoting and inhibiting inflammation among different groups of T cells, such as Th1, Th2, Tregs, and Th17. Tregs play an essential immunomodulatory role in reducing its immunopathological reaction[2, 19, 39, 40]. To illustrate the effect and possible mechanism of unstable Tregs on *Schistosoma japonicum* infection, and to study whether USP21 plays a new role in controlling the stability of Tregs, we used C57BL/6 mice model with the conditional knockout USP21 in Tregs, which was infected *Schistosoma japonicum* and was often used as model high-vs.low-pathology strains [41].

Schistosomes infection can be divided into three stages and can present in different clinical symptoms and immune mechanisms: acute infection, active infection, and late chronic infection [42]. Our study found that after *Schistosoma japonicum* infection, the number of coups and adults recovered from the KO mice was much larger than from the WT mice, and the same trend was noted for the number of the eggs. At the same time, we noted that there was no gender difference in adults between the two groups. It is known that Schistosoma depended on host signals to keep correct development and maturation, such as TGF-α and TNF-α [43]. We speculated that the changes in the immune microenvironment in the host might affect the binding of *Schistosoma japonicum*. However, there was no apparent difference in the liver and spleen between KO and WT mice with *Schistosoma japonicum*. Considering that the protection of *Schistosoma japonicum* infection was mainly based on the clearance of *Schistosoma japonicum* in the early infection stage, it also seemed that USP21^-/-^ in Tregs might inhibit the lethal abilities of *Schistosoma Japonicum*. These two theories need to be further evaluated in the future.

Th1 immune response mainly in the early stage of *Schistosoma Japonicum* infection was to eliminate adults, and Th2 immune response was mainly to anti-eggs in the later stage[44]. Liver egg granuloma was the primary immunopathological reaction, the severity of the symptom of which was usually related to the intensity of infection [42]. We found that the diameter of egg granuloma in KO mice was smaller than that in WT mice at six weeks, and the infiltration of inflammatory-related substances such as neutrophils was less and lighter. The degree of hepatic fibrosis in KO mice was also lower than that in WT mice by RT-PCR, Western blot, and Masson Staining. The immune response of T cells fluctuates at different levels during the infection of *Schistosoma japonicum*, Th1, and T-reg are co-dominant in transcription level, Th1 and Th2 are co-dominant in protein level [45]. These results suggested that unstable Treg cells may reduce the pathological damage of the liver through some mechanism.

Cytokines produced by egg antigen-stimulated liver T cell populations from infected mice are faithful correlates of immunopathology. High-pathology strains sustain a proinflammatory Th1 and Th17(IL-17) cell-polarized response alongside the Th2 response [41], whereas an initial short-lived elevation in IFN-γ-producing Th1 cells gives way to a mostly host-protective Th2(IL-4, IL-5, IL-13)-dominated environment [46]. We found that the content of IFN-γ and IL-4 in KO mice was higher than that in WT mice, while IL-10, IL-17, IL-23 and IL-9 was lower than that in the WT group. With different durations of infection, the trend of cytokines changes in the serum of two groups was about the same. These results are partly consistent with the previous findings, and once again verify that USP21 knock-out will shift Treg to Th1-like cells[26]. IFN-γ content in the KO group was much higher than that in the WT group, and IFN-γ in the later stage of *Schistosoma japonicum* infection was maintained stably with IL-4 increasing. However, it was different from those mentioned above that IL-4 content was higher in KO mice than that in WT mice. The unstable Treg might have produced different differentiation effects because of the different microenvironments of the schistosome infection, and series of T-cell activation and long-term stimulation of chronic inflammatory factors could have low reactivity [26, 47, 48]. This coincidence was in line with the weakening effect of other secreted factors in addition to IFN-γ and IL-4. It was also speculated that Th1 and Th2 are the main immunity types of the Schistosomiasis. According to the previous reports, Th17 might play an essential role in the formation and growth of the liver immunopathology and the egg granuloma caused by Schistosoma adults, and the development of severe Schistosomiasis was related to the high level of IL-17[49, 50]. The results of this study also confirmed that IL-17 content in the KO group was less than that in the WT group, and the pathological sections of the liver revealed, as shown in Fig.2B. IL-23 was one of the driving factors to produce IL-17. We also observed that IL-17 was related to IL-23, both of which were lower in the KO group than that in the WT group, and both were down-regulated.

The liver fibrosis and spleen diseases caused by *Schistosoma japonicum* were associated with high FOXP3^+^Tregs in the blood[18]. FOXP3 played an important role in regulating the differentiation, development, and function of Tregs, and the three FOXP3, GATA3 and USP21 are closely related to Tregs. To clarify the relation between USP21 and FOXP3^+^Tregs, our results of CD4^+^CD25^+^FOXP3^+^Tregs were consistent with previous studies after spleen cell isolation despite some differences. The percentage of FOXP3^+^Tregs in the KO group was lower than that in the WT group, and the percentage of CD4^+^T cells was higher than that in the WT group. The results indicated that the deletion of USP21 made the expression of FOXP3 unstable and led to Spontaneous activation and increasing of T cells. FOXP3^+^Tregs were reduced in the KO group because of the easy degradation of unstable FOXP3. Another evidence supported that FOXP3 instability promoted the production of T-like Treg cells.

Soluble egg antigen (SEA) and adult worm antigen (SWA) are the main soluble proteins related to adaptive immune response induced by *Schistosoma japonicum* infection. Related studies have shown that a high level of anti-Schistosoma IgG was associated with increased susceptibility to parasites. These studies have demonstrated a positive correlation between anti-Schistosoma IgG, especially IgG4 response and severe Schistosomiasis[51, 52]. In this study, the levels of SEA and SWA IgG in USP21 knock-out mice infected with *Schistosoma japonicum* were higher than those in the WT group while IgM levels were the same or lower. This finding suggested that unstable Treg cells improved the immune response to Schistosoma and reduced the initial immune response and lethality to Schistosoma.

Worms live in the host for a long time, and there is growing evidence that they can manipulate the host’s immune system through the host’s immunomodulatory. The immunomodulatory network is activated after chronic worm infection and can even cross-regulate the non-associated inflammatory response of the host. It has been proven that FOXP3^+^ and Tregs were essential parts of this regulatory network, especially for the immunosuppression of chronic Schistosomiasis, which was characterized by the independent down-regulation of IL-10 produced by Th2 cytokines [53-55]. Therefore, it is suggested that unstable Tregs could also be used by the parasite to avoid the immune system of the host, which may provide a new research basis for the immunoregulation between the host and the parasite, and is of considerable significance to the treatment of the chronic inflammatory disease.

In conclusion, the significant heterogeneity of the host’s immune response to schistosome infection, involving different APC, T-cell subsets and cytokines, led to different immunopathology levels. Our study still found that unstable Tregs in mice infected with *Schistosoma japonicum* might not only benefit the host from the excessive inflammatory response but also regulate the living microenvironment of parasites, both of which achieved a new balance. There were still some limitations in this study because many factors, such as the subsequent survival status of mice, were not considered. However, this paper could provide a theoretical basis to study the regulation of USP21 further on different immune cell groups induced by *Schistosoma Japonicum*, and provide a new possible target for schistosomiasis treatment (anti-*Schistosoma japonicum*) in the future.

## Materials and Methods

### Ethics statement

We carried out all animal experiments in strict accordance with the Laboratory Animal Regulation (1988.11.1) and made every effort to minimize the suffering. The IACUC of National Institution of Parasitic Diseases of Chinese Center for Diseases Control and Prevention and Control approved all animal procedures for the use of experimental animals (License No: NJMU 07-0137).

### Experimental animals, parasites, and the establishment of infection models

Two kinds of female mice, FOXP3^Cre^ (WT) and USP21^fl/fl^FOXP3^Cre^ (KO), were provided by Shanghai Institute of Bioscience, Chinese Academy of Sciences. *Schistosoma japonicum* cercariae escaped from naturally infected snails from the National Institute of Parasitic Diseases of Chinese Center for Diseases Control and Prevention (Shanghai, China). Each of the two kinds mice was divided into a normal control group (NC/Uninfected) (n =2) and infection group (INF) (n =10 ± 2) for 6-8 weeks under specific pathogen-free conditions, then the mice of INF group were infected with 25 ± 1 cercariae via abdomen and killed all at 6-7weeks after infection, including mice of Uninfected group. Serum, stored at −80°C and used for ELISA, was collected from mice blood at different times, respectively before challenge infection, 20 days and 30days after infection, and 42 days after challenge infection with eyeball enucleation. The animal experiments were carried out randomly by blind strategies, strictly per the Laboratory Animal Regulation, and we made every effort to minimize pain. The model of mice infected with the *Schistosoma japonicum* was established by repeating three times.

### Collection of *Schistosoma japonicum* worms and weight of the spleen and liver of the infected mice

The mice, 42 days after challenge infection, were killed by the cervical dislocated method. Worms were collected from the portal vein system by cardiac infusion of saline, and the number of male and female adult worms and hugs was determined. The spleen and liver of the mice were weighed and photographed.

### HE and Masson staining

The same site of each infected mouse was fixed in 4% paraformaldehyde and embedded in the paraffin block. We stained a slice (5μm thick) with hematoxylin-eosin and performed the analysis with a microscope (magnification ×100). To evaluate the pathological changes and the number of eggs among groups, the area of liver single egg granuloma (n ≥ 5 per mouse) was measured by software, and the average number of eggs was counted by a random selection of visual field (n ≥ 5 per mouse) at magnification × 40. Similarly, the same site of the liver was stained with Masson. In order to evaluate the degree of hepatic fibrosis between the two groups, we measured and analyzed the ratio of collagen area (n ≥ 5 per mouse) using appropriate software.

### Isolation of Peripheral Blood lymphocytes (PBMC) and Culture in vitro

0.5-1ml blood was taken from mice 42 days after infection by eyeball enucleation, serum and plasma were separated, and serum was stored at −80°C to be studied. We mixed the plasma with sterilized PBS in the proportion of 1:1, and the mixture was gently reversed back and forth for about ten times and gently added the same volume of lymphocyte isolation to form stratification. After centrifugation, the study team took the middle layer to add red blood cell lysate to decompose red cells, centrifuge again to get PBMC. The PBMC were resuspended with a complete culture medium containing 5 μg /ml RMPI1640. The cell density under the microscope was adjusted to 2×10^5^-10^6^/ml, then divided into culture plate or dish, and cultured at 37°C, 5% CO_2_ for 72 h under sterile conditions. We then stored the culture supernatant after centrifugation at −80°C.

### Cell Isolation of Spleen and Culture in vitro

The mice, 42 days after infection, were killed by the cervical dislocated method to isolate the spleen. The spleen passed through a 100-μm cell strainer to get tissue homogenate and was added red blood cell lysate to decompose red cells repeatedly until there were no red blood cells. Then a complete culture medium containing 5 μg /ml RMPI1640 was added to resuspend cells. The cell density under the microscope was adjusted to 2×10^5^-10^6^/ml, then divided into culture plate or dish, and cultured at 37°C, 5%CO_2_ for 72h under sterile conditions. The culture fluids centrifuged to remove supernatant were stored at-80°C, The single-cell suspension was detected by flow cytometry.

### Flow Cytometry

For cell-surface marker and Treg cells analysis, CD4^+^ T cells were isolated from 5×10^5^-10^6^/ml single-cell suspension using the CD4 (L3T4) MicroBeads, which was permeabilized in PBS with 2% fetal bovine serum, and then were stained on the surface. The cell membrane was destroyed, and the nuclear transcription factor FOXP3 was stained according to the instructions of the manufacturer. Compensation was performed with BD LSRFortessa (BD Biosciences). The data were acquired and analyzed with FlowJo (Tree Star) software.

### RNA extraction and RT-PCR

We extracted the total RNA from 30 mg liver and spleen with Trizol, based on the manual. The quality and quantity of RNA were then evaluated by ultra-micro-nucleic acid protease analyzer. Total RNA (1μg) was reverse transcribed with RevertAid First Strand cDNA Synthesis Kit (K1622, Thermo Fisher Scientific). The expression of USP21, IL-10, IL-17, FOXP3 in liver and spleen, and α-SMA, collagenI, collagen III in the liver were detected by iTaq™ Universal SYBR® Green super mixture (1725121, bio-rad) and fluorescence quantitative PCR 7500 system (applied biosystems, USA), and quantitative analysis was carried out by 2^-ΔΔCt^ method with glyceraldehyde-3-phosphate dehydrogenase (GAPDH) as control. The PCR cycling conditions were as follows: 40 cycles of 95°C for 30s; 95°C for 5s; 60°C for 34s; and then 95°C for 15s; 60°C for 60s; 95°C for 15s; and 60°C for 60s. All primers were blasted to NCBI to ensure their specificity. All primers were showed as Table 1

**Table 1.**
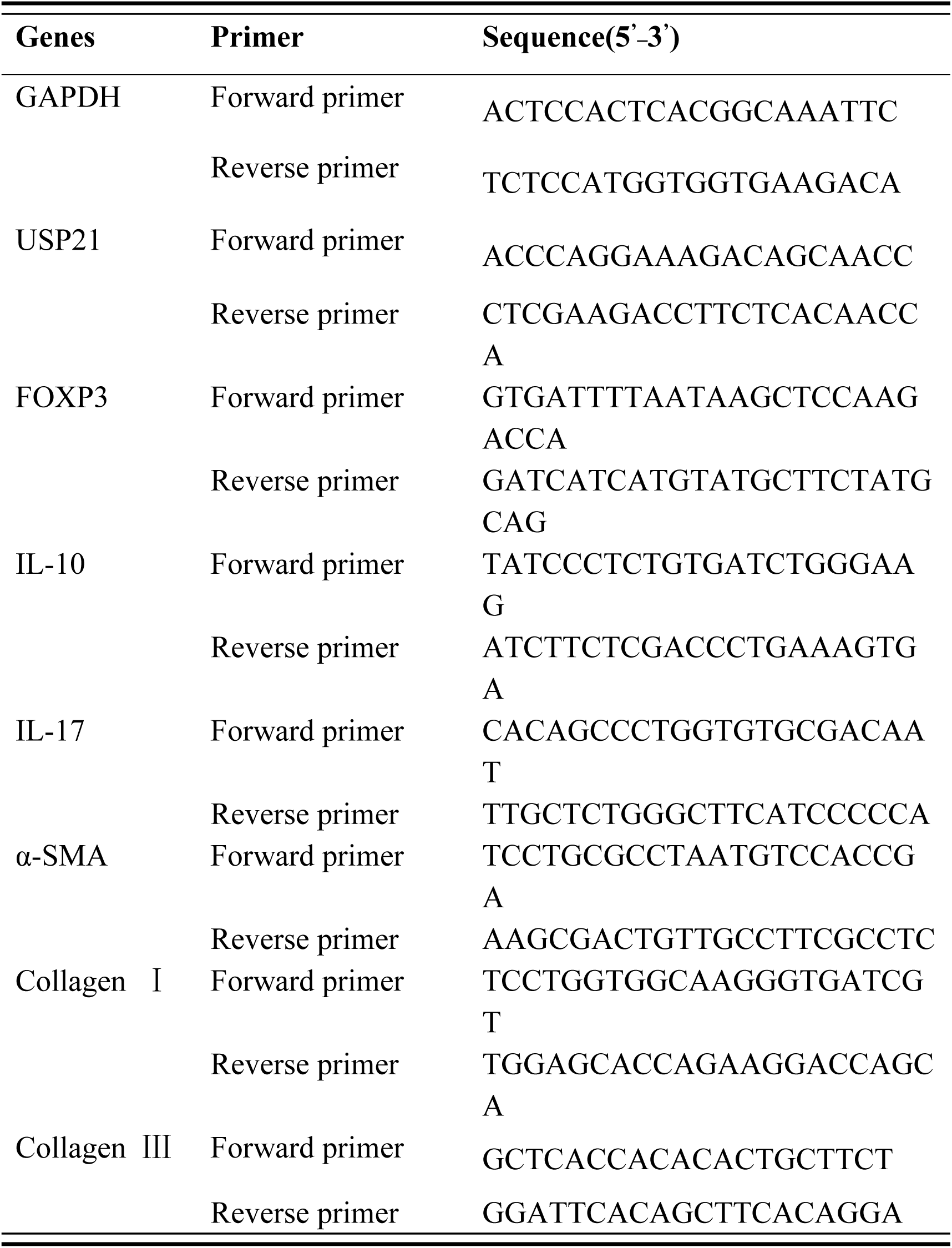
Primers used for mRNA analysis.

### Western Blotting

30-50mg liver from each mouse was lysed on ice for 3 to 4 hours by the cold ripa lytic buffer(1-1.2ml) containing cocktail protease inhibitor (B14001, bimake). After centrifuging 15min at 14000g/min, the supernatant was moved to another tube to detected the total protein concentration by BCA protein detection kit (P0010, Biyuntian). The protein was boiled for 3 min and loaded onto 10% polyacrylamide gel, electrophoresed, and transferred to PVDF membrane (MilliporeMA, USA). The proteins were separated by SDS–gel electrophoresis. Protein bands detected by incubation for12-16h at 4°C with polyclonal primary antibodies to GAPDH(sc-32233, Santa Cruz), USP21(ab171028, abcam), α-SMA(#19245T, Cell Signaling Technology), Collagen I (BA0325, Boster), Collagen III(sc-271249, Santa Cruz) followed by blotting with secondary antibody HRP-Goat anti-Rabbit IgG (ab97080, Abcam), HRP-Rabbit anti-Mouse IgG(ab6728, Abcam). Immune complexes were visualized with WesternBright^™^ECL substrate (K-12045-D10, Advansta) and the luminescent signal recorded in the chemiluminescence imaging system (Bio-Rad, USA) to detect the expression of proteins. Quantitative analysis of the expression of USP21, α-SMA, Collagen I, Collagen III was carried out by using ImageJ (NIH, Bethesda, USA).

### ELISA

The ELISA assay has been described previously[56]. Briefly, for the *S.j*-SEA-, and *S.j*-SWAP-ELISAs, all recombinant antigens were diluted to a final concentration of 1 μg/mL with coating buffer; for the *S.j*-SEA-and *S.j*-SWAP-ELISAs, 50ng of each antigen was mixed per well at 4°C overnight with 100 μL added per well. After blocking by blocking buffer (1% BAS in PBST) at 37°C for one hour. Then serum samples diluted at 1:250 with blocking buffer were added (100 μL/well) and incubated for one hour at 37°C. HRP-Goat anti-Mouse IgM (ab97230, abcam)and HRP-Rabbit anti-Mouse IgG(ab6728, abcam) were used as the secondary antibody (1:20,000, 100 μL/well), and samples were incubated for one hour at 37°C. Streptavidin-HRP (BD Pharmingen, CA, USA) (1:10,000) was then applied to each well (100 μL/well). PBST washes were applied five times after each step, 2 min between each wash. Reactions were developed using TMB as substrate (100 μL/well) for 5 min and stopped using 2 M sodium hydroxide (50 μL/well). Optical density (OD) values were read at 450 nm using a microplate reader, and all tests were run in duplicate on each test plate.

### Cytokine detection (Multiplex Fluorescent Microsphere Immunoassay)

The mouse serum and cell culture samples were prepared according to the manufacturer’s instructions. 50µl pre-mixed beads were added to each of 96 wells and then were incubated with the samples. Detecting antibodies (IFN-γ, IL-4, IL-10, IL-17A, IL-23, IL-9) were added after washing, followed by adding SA-PE after washing again. The 96-wells clean-dried plate was put into Bio-Plex 200 to detect. The standard curve was fitted by a five-parameter non-linear regression method, and the concentration was calculated. The results included the label, the median of the fluorescence intensity and the concentration.

### Statistical analyses

Statistical comparisons were performed with Prism 7.0 (GraphPad Prism.) and SPSS18.0, using *t*-test for comparisons of two datasets, and ANOVA for multiple comparisons. All the data were set for 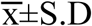 and tests were considered statistically significant at *p*≤0.05.

## Acknowledgment

We thank Dr. Adiele Onyeze for thorough reviews that greatly improved the manuscript.

## References

1. Ross AG, Bartley PB, Sleigh AC, Olds GR, Li Y, Williams GM, et al. (2002) Schistosomiasis. N Engl J Med 346:1212–20. pmid: 11961151

2. Wilson MS, Mentink-Kane MM, Pesce JT, Ramalingam TR, Thompson R, Wynn TA (2007) Immunopathology of schistosomiasis. Immunology and Cell Biology 85:148–54. pmid: WOS:000245102500011

3. Yazdanbakhsh M, Sacks DL (2010) Why does immunity to parasites take so long to develop? Nat Rev Immunol 10:80–1. pmid:WOS:000273876200001

4. Allen JE, Maizels RM (2011) Diversity and dialogue in immunity to helminths. Nat Rev Immunol 11:375–88. pmid: 21610741

5. Nausch N, Bourke CD, Appleby LJ, Rujeni N, Lantz O, Trottein F, et al. (2012) Proportions of CD4+ memory T cells are altered in individuals chronically infected with Schistosoma haematobium. Sci Rep 2:472. pmid: 22737405

6. McSorley HJ, Maizels RM (2012) Helminth infections and host immune regulation. Clin Microbiol Rev 25:585–608. pmid: 23034321

7. Yang Q, Qu J, Jin C, Feng Y, Xie S, Zhu J, et al. (2019) Schistosoma japonicum Infection Promotes the Response of Tfh Cells Through Down-Regulation of Caspase-3-Mediating Apoptosis. Front Immunol 10:2154. pmid: 31572373

8. Johnson PT, Hoverman JT (2012) Parasite diversity and coinfection determine pathogen infection success and host fitness. Proc Natl Acad Sci U S A 109:9006–11. pmid: 22615371

9. Riner DK, Ferragine CE, Maynard SK, Davies SJ (2013) Regulation of innate responses during pre-patent schistosome infection provides an immune environment permissive for parasite development. PLoS Pathog 9:e1003708. pmid: 24130499

10. Zhan T, Ma H, Jiang S, Zhong Z, Wang X, Li C, et al. (2019) Interleukin-9 blockage reduces early hepatic granuloma formation and fibrosis during Schistosoma japonicum infection in mice. Immunology 158:296–303. pmid: 31436861

11. Liu J, Zhu L, Wang J, Qiu L, Chen Y, Davis RE, et al. (2019) Schistosoma japonicum extracellular vesicle miRNA cargo regulates host macrophage functions facilitating parasitism. PLoS Pathog 15:e1007817. pmid: 31163079

12. Colley DG, Secor WE (2014) Immunology of human schistosomiasis. Parasite Immunol 36:347–57. pmid: 25142505

13. Burns LA, Maroof A, Marshall D, Steel KJA, Lalnunhlimi S, Cole S, et al. (2019) Presence, function, and regulation of IL-17F-expressing human CD4(+) T cells. Eur J Immunol. pmid: 31850514

14. Razawy W, Asmawidjaja PS, Mus AM, Salioska N, Davelaar N, Kops N, et al. (2019) CD4(+) CCR6(+) T cells, but not gammadelta T cells, are important for the IL-23R-dependent progression of antigen-induced inflammatory arthritis in mice. Eur J Immunol. pmid: 31778214

15. Chen X, Yang X, Li Y, Zhu J, Zhou S, Xu Z, et al. (2014) Follicular helper T cells promote liver pathology in mice during Schistosoma japonicum infection. PLoS Pathog 10:e1004097. pmid: 24788758

16. Zaretsky AG, Taylor JJ, King IL, Marshall FA, Mohrs M, Pearce EJ (2009) T follicular helper cells differentiate from Th2 cells in response to helminth antigens. J Exp Med 206:991–9. pmid: WOS:000266010000006

17. Zhang YM, Wang YJ, Jiang YY, Pan W, Liu H, Yin JH, et al. (2016) T follicular helper cells in patients with acute schistosomiasis. Parasite Vector 9. pmid: WOS:000377338600001

18. Romano A, Hou X, Sertorio M, Dessein H, Cabantous S, Oliveira P, et al. (2016) FOXP3+ Regulatory T Cells in Hepatic Fibrosis and Splenomegaly Caused by Schistosoma japonicum: The Spleen May Be a Major Source of Tregs in Subjects with Splenomegaly. PLoS Negl Trop Dis 10:e0004306. pmid: 26731721

19. Khattri R, Cox T, Yasayko SA, Ramsdell F (2003) An essential role for Scurfin in CD4(+)CD25(+) T regulatory cells. Nat Immunol 4:337–42. pmid: WOS:000181987700012

20. Fontenot JD, Gavin MA, Rudensky AY (2003) Foxp3 programs the development and function of CD4+CD25+ regulatory T cells. Nat Immunol 4:330–6. pmid: 12612578

21. Hori S, Nomura T, Sakaguchi S (2003) Control of regulatory T cell development by the transcription factor Foxp3. Science 299:1057–61. pmid: 12522256

22. Taylor MD, van der Werf N, Maizels RM (2012) T cells in helminth infection: the regulators and the regulated. Trends Immunol 33:181–9. pmid: 22398370

23. Turner JD, Jenkins GR, Hogg KG, Aynsley SA, Paveley RA, Cook PC, et al. (2011) CD4+CD25+ regulatory cells contribute to the regulation of colonic Th2 granulomatous pathology caused by schistosome infection. PLoS Negl Trop Dis 5:e1269. pmid: 21858239

24. Zhou X, Bailey-Bucktrout SL, Jeker LT, Penaranda C, Martinez-Llordella M, Ashby M, et al. (2009) Instability of the transcription factor Foxp3 leads to the generation of pathogenic memory T cells in vivo. Nat Immunol 10:1000–7. pmid: 19633673

25. Samstein RM, Arvey A, Josefowicz SZ, Peng X, Reynolds A, Sandstrom R, et al. (2012) Foxp3 exploits a pre-existent enhancer landscape for regulatory T cell lineage specification. Cell 151:153–66. pmid: 23021222

26. Li Y, Lu Y, Wang S, Han Z, Zhu F, Ni Y, et al. (2016) USP21 prevents the generation of T-helper-1-like Treg cells. Nat Commun 7:13559. pmid: 27857073

27. Wan YY, Flavell RA (2007) Regulatory T-cell functions are subverted and converted owing to attenuated Foxp3 expression. Nature 445:766–70. pmid: 17220876

28. Levescot A, Nelson-Maney N, Morris A, Bouyer RG, Lee P, Nigrovic P (2017) Autoimmune Arthritis in IL-1 Receptor Antagonist-Deficient Mice Is Associated with a Pathogenic Conversion of Foxp3+Regulatory T Cells into Th17 Cells. Arthritis Rheumatol 69. pmid: WOS:000411824106216

29. Dominguez-Villar M, Baecher-Allan CM, Hafler DA (2011) Identification of T helper type 1-like, Foxp3(+) regulatory T cells in human autoimmune disease. Nat Med 17:673–5. pmid: WOS:000291308800026

30. Bailey-Bucktrout SL, Martinez-Llordella M, Zhou X, Anthony B, Rosenthal W, Luche H, et al. (2013) Self-antigen-driven activation induces instability of regulatory T cells during an inflammatory autoimmune response. Immunity 39:949–62. pmid: 24238343

31. Zhang J, Chen C, Hou XX, Gao YY, Lin F, Yang J, et al. (2013) Identification of the E3 Deubiquitinase Ubiquitin-specific Peptidase 21 (USP21) as a Positive Regulator of the Transcription Factor GATA3. J Biol Chem 288:9373–82. pmid: WOS:000316862200054

32. Fan YH, Mao RF, Yu Y, Liu SF, Shi ZC, Cheng J, et al. (2014) USP21 negatively regulates antiviral response by acting as a RIG-I deubiquitinase. J Exp Med 211:313–28. pmid: WOS:000331281600011

33. Chen Y, Wang L, Jin J, Luan Y, Chen C, Li Y, et al. (2017) p38 inhibition provides anti-DNA virus immunity by regulation of USP21 phosphorylation and STING activation. J Exp Med 214:991–1010. pmid: 28254948

34. Nakagawa T, Kajitani T, Togo S, Masuko N, Ohdan H, Hishikawa Y, et al. (2008) Deubiquitylation of histone H2A activates transcriptional initiation via trans-histone cross-talk with H3K4 di- and trimethylation. Genes Dev 22:37–49. pmid: 18172164

35. Tao LQ, Chen C, Song HH, Piccioni M, Shi G, Li B (2014) Deubiquitination and stabilization of IL-33 by USP21. Int J Clin Exp Patho 7:4930–7. pmid: WOS:000345120900036

36. Arceci A, Bonacci T, Wang X, Stewart K, Damrauer JS, Hoadley KA, et al. (2019) FOXM1 Deubiquitination by USP21 Regulates Cell Cycle Progression and Paclitaxel Sensitivity in Basal-like Breast Cancer. Cell Rep 26:3076–86 e6. pmid: 30865895

37. Liu FW, Fu Q, Li YP, Zhang K, Tang MY, Jiang W, et al. (2019) USP21 modulates Goosecoid function through deubiquitination. Bioscience Rep 39. pmid: WOS:000475404500001

38. Pannu J, Belle JI, Forster M, Duerr CU, Shen SY, Kane L, et al. (2015) Ubiquitin Specific Protease 21 Is Dispensable for Normal Development, Hematopoiesis and Lymphocyte Differentiation. Plos One 10. pmid: WOS:000350682600061

39. Lacorcia M, Prazeres da Costa CU (2018) Maternal Schistosomiasis: Immunomodulatory Effects With Lasting Impact on Allergy and Vaccine Responses. Front Immunol 9:2960. pmid: 30619318

40. Tang CL, Zhang RH, Liu ZM, Jin H, He L (2019) Effect of regulatory T cells on the efficacy of the fatty acid-binding protein vaccine against Schistosoma japonicum. Parasitol Res 118:559–66. pmid: 30607606

41. Larkin BM, Smith PM, Ponichtera HE, Shainheit MG, Rutitzky LI, Stadecker MJ (2012) Induction and regulation of pathogenic Th17 cell responses in schistosomiasis. Semin Immunopathol 34:873–88. pmid: 23096253

42. McManus DP, Dunne DW, Sacko M, Utzinger J, Vennervald BJ, Zhou XN (2018) Schistosomiasis. Nat Rev Dis Primers 4:13. pmid: 30093684

43. Amiri P, Locksley RM, Parslow TG, Sadick M, Rector E, Ritter D, et al. (1992) Tumour necrosis factor alpha restores granulomas and induces parasite egg-laying in schistosome-infected SCID mice. Nature 356:604–7. pmid: 1560843

44. Kalantari P, Bunnell SC, Stadecker MJ (2019) The C-type Lectin Receptor-Driven, Th17 Cell-Mediated Severe Pathology in Schistosomiasis: Not All Immune Responses to Helminth Parasites Are Th2 Dominated. Front Immunol 10:26. pmid: 30761125

45. Farwa A, He C, Xia L, Zhou H (2018) Immune modulation of Th1, Th2, and T-reg transcriptional factors differing from cytokine levels in Schistosoma japonicum infection. Parasitol Res 117:115–26. pmid: 29188369

46. Pearce EJ, Caspar P, Grzych JM, Lewis FA, Sher A (1991) Downregulation of Th1 cytokine production accompanies induction of Th2 responses by a parasitic helminth, Schistosoma mansoni. J Exp Med 173:159–66. pmid: 1824635

47. Stadecker MJ, Asahi H, Finger E, Hernandez HJ, Rutitzky LI, Sun J (2004) The immunobiology of Th1 polarization in high-pathology schistosomiasis. Immunol Rev 201:168–79. pmid: 15361240

48. El-Ahwany E, Bauiomy IR, Nagy F, Zalat R, Mahmoud O, Zada S (2012) T regulatory cell responses to immunization with a soluble egg antigen in Schistosoma mansoni-infected mice. Korean J Parasitol 50:29–35. pmid: 22451731

49. Rutitzky LI, Stadecker MJ (2006) CD4 T cells producing pro-inflammatory interleukin-17 mediate high pathology in schistosomiasis. Mem Inst Oswaldo Cruz 101 Suppl 1:327–30. pmid: 17308791

50. Wen XY, He L, Chi Y, Zhou S, Hoellwarth J, Zhang C, et al. (2011) Dynamics of Th17 Cells and Their Role in Schistosoma japonicum Infection in C57BL/6 Mice. Plos Neglect Trop D 5. pmid: WOS:000298134000027

51. Yang Q, Qiu HN, Xie HY, Qi YW, Cha HF, Qu JL, et al. (2017) A Schistosoma japonicum Infection Promotes the Expansion of Myeloid-Derived Suppressor Cells by Activating the JAK/STAT3 Pathway. Journal of Immunology 198:4716–27. pmid: WOS:000405271300018

52. Negrao-Correa D, Fittipaldi JF, Lambertucci JR, Teixeira MM, Antunes CMD, Carneiro M (2014) Association of Schistosoma mansoni-Specific IgG and IgE Antibody Production and Clinical Schistosomiasis Status in a Rural Area of Minas Gerais, Brazil. Plos One 9. pmid: WOS:000336971300043

53. Wynn TA, Cheever AW, Williams ME, Hieny S, Caspar P, Kuhn R, et al. (1998) IL-10 regulates liver pathology in acute murine Schistosomiasis mansoni but is not required for immune down-modulation of chronic disease. J Immunol 160:4473–80. pmid: 9574553

54. Maizels RM, Balic A, Gomez-Escobar N, Nair M, Taylor MD, Allen JE (2004) Helminth parasites - masters of regulation. Immunological Reviews 201:89–116. pmid: WOS:000223819600008

55. Baumgart M, Tompkins F, Leng J, Hesse M (2006) Naturally occurring CD4+Foxp3+ regulatory T cells are an essential, IL-10-independent part of the immunoregulatory network in Schistosoma mansoni egg-induced inflammation. J Immunol 176:5374-87. Epub 2006/04/20. pmid: 16622005

56. Cai P, Weerakoon KG, Mu Y, Olveda DU, Piao X, Liu S, et al. (2017) A Parallel Comparison of Antigen Candidates for Development of an Optimized Serological Diagnosis of Schistosomiasis Japonica in the Philippines. EBioMedicine 24:237-46. Epub 2017/09/26. pmid: 28943229

